# Application of Engineered NK-92 Cell Extracellular Vesicles in the Treatment of Systemic Lupus Erythematosus

**DOI:** 10.64898/2026.04.14.718139

**Authors:** Yu Sun, Zeyu Tang, Mengting Guo, Zimeng Zhai, Zixian Wu, Xia Wang, Fang Li, Weiling An, Xiaowei Dou

## Abstract

**Introduction:** Systemic Lupus Erythematosus (SLE) is a chronic autoimmune disease characterized by loss of immune tolerance, autoantibody production, and multi-organ damage. Current therapies, including glucocorticoids and CAR-T/CAR-NK cell therapies, are limited by adverse effects, high cost, and safety concerns.

**Objectives:** To develop engineered NK-92 cell-derived extracellular vesicles displaying CD19 single-chain variable fragment (V-CD19-Exo) and evaluate their therapeutic efficacy in an MRL/lpr mouse model of SLE.

**Methods:** The CD19scFv-LAMP-2B fusion construct was stably expressed in NK-92 cells via lentiviral transduction. Extracellular vesicles were isolated by differential centrifugation and characterized by NTA, TEM, and Western Blot. In vivo efficacy was assessed in MRL/lpr mice through B cell depletion analysis, renal function monitoring, cytokine profiling, autoantibody detection, and survival observation.

**Results:** V-CD19-Exo significantly reduced splenic CD19□CD20□ B cells from 10.53% to 1.51% (p < 0.0001). Treatment attenuated proteinuria, ameliorated lupus nephritis pathology, reversed splenomegaly, and downregulated serum IgE, IL-17A, IFN-γ, anti-dsDNA, and ANA levels. Notably, V-CD19-Exo improved survival to approximately 80% compared to 25% in untreated controls.

**Conclusion:** Engineered NK-92 cell-derived extracellular vesicles represent a novel, safe, and effective cell-free therapeutic strategy for SLE, offering advantages over conventional cell therapies including lower immunogenicity, scalable production, and no requirement for lymphodepletion.

## 1. Introduction

Systemic Lupus Erythematosus (SLE) is a systemic inflammatory disease characterized by autoimmune reactions, primarily featuring loss of immune tolerance, massive production of autoantibodies, and deposition of immune complexes in multiple organs. The global prevalence is approximately 43.7 cases per 100,000 people, with higher incidence rates in women of childbearing age and Asian populations, showing significant gender and regional disparities [1, 2]. SLE manifests with diverse clinical symptoms, potentially affecting multiple organs including the skin, joints, kidneys, and nervous system. Among these, Lupus Nephritis (LN), as one of the most common end-organ damages, is often the main cause of renal failure and death, with approximately 40%–60% of SLE patients showing renal involvement at the early stage of the disease, significantly impacting prognosis [3–5]. The pathogenesis of SLE remains unclear, involving complex interactions among genetics, epigenetics, environmental factors, and immune system dysregulation. The core mechanism involves the abnormal activation of B cells; autoreactive B cells not only produce large amounts of autoantibodies but also promote T cell activation through antigen presentation and cytokine secretion, forming an autoimmune loop [6–8]. B Lymphocyte Stimulator (BLyS) and A PRoliferation-Inducing Ligand (APRIL) promote the survival and differentiation of autoreactive B cells by binding to BR3, TACI, and BCMA receptors on the B cell surface [9]. CD19, as a specific surface marker for B cells, is widely expressed from pro-B cells to mature B cells and plays a crucial role in B cell activation, proliferation, and differentiation. Therefore, B cell-related targets (CD20, CD19, CD40L, BAFF, etc.) have become focal points in drug development in recent years.

Current treatment for SLE primarily relies on glucocorticoids and immunosuppressants. Although these can control the disease, long-term use may be accompanied by severe adverse effects such as infections, metabolic disorders, and organ toxicity [4]. In recent years, targeted therapeutic strategies have made some progress. For instance, monoclonal antibodies targeting B-cell activating factor (BAFF/BLyS) or CD20 have shown significant efficacy in alleviating symptoms for some patients. However, clinical practice and studies indicate that their overall efficacy is limited by factors such as uneven depletion [10–12].

Cell therapy, mainly CAR-T cell therapy, is not only applied in treating hematological malignancies, particularly B-cell leukemia and lymphoma [13, 14], but also offers a highly promising potential technology for treating autoimmune diseases. Multiple clinical studies have confirmed that CD19-targeted CAR-T cell therapy can efficiently eliminate mature B cells and precursor B cells in vivo, leading to significant improvement in clinical symptoms and decreased autoantibody levels in patients with refractory SLE and psoriasis [15–17].

Gao et al. successfully applied allogeneic CD19-targeted CAR-NK technology in a First-in-Human clinical study for SLE. Among 9 patients followed for over 1 year, 6 patients (67%) achieved complete DORIS remission and reached a lupus low disease activity state, with no recurrence observed in any patient [18]. Wang et al. reported on iPSC-derived CAR-NK cell therapy for systemic sclerosis [19].

However, CAR-T and CAR-NK cell therapies face limitations including high cost, lengthy preparation time, requirement for lymphodepletion before treatment, and risks such as cytokine release syndrome (CRS). Allogeneic CAR-NK cell therapy also encounters challenges like low expansion efficiency, insufficient persistence in vivo, and high costs [15, 20], restricting the application of CAR-T/CAR-NK in treating autoimmune diseases. Therefore, continuing to explore safer, more precise, and more accessible next-generation therapies has become a crucial research direction.

Extracellular vesicles are nanoscale vesicles (approximately 30–150 nm in diameter) secreted by various cells and present in body fluids. Naturally carrying bioactive molecules such as proteins, nucleic acids, and lipids, they are important mediators of intercellular communication, possessing low immunogenicity, high stability, excellent nanomaterial properties, and tissue penetration capability [21–30]. Numerous studies have confirmed that extracellular vesicles inherit the functions of their parent cells. For example, mesenchymal stem cell-derived exosomes (MSC-exos) show significant potential in regulating immune responses and promoting tissue repair. By delivering functional molecules like miRNAs, they regulate macrophage polarization, promote regulatory T cell (Treg) proliferation, and inhibit aberrant immune responses [28, 31]. Exosomes derived from Natural Killer cells or engineered CAR-NK cells have demonstrated therapeutic effects in melanoma and breast cancer therapy, retaining tumor cell killing functions. More importantly, they offer higher safety, a wide source, and are easy to preserve and transport [32–35].

However, naturally produced extracellular vesicles have limitations such as low yield and poor targeting. Engineered modification strategies, including parental cell gene editing, membrane surface modification, and cargo loading, can address these challenges [36–40]. NK-92 cells, as a natural immune cell line with strong cytotoxicity and easy expandability, have been successfully used in numerous clinical trials for cancer treatment (e.g., patients with recurrent HER2-positive glioblastoma receiving intracranial injection of CAR-NK-92 cells; patients with refractory/relapsed AML receiving NK-92 cell infusions) [41, 42]. Extracellular vesicles derived from NK-92 cells also inherit the killing activity and immunomodulatory capacity of NK cells, showing promise for cancer therapy with good clinical translation potential [43–46]. NK-92 exosomes have demonstrated cytotoxicity against leukemia HL-60 cells [27].

Based on the above background, we utilized NK-92 cell-derived extracellular vesicles to construct an engineered nanovesicle platform combining targeting ability and immunomodulatory functions. We fused the CD19 single-chain variable fragment (CD19scFv) with the N-terminus of Lysosome-Associated Membrane Protein 2B (LAMP-2B), constructed a recombinant plasmid, and transfected it into NK-92 cells, thereby displaying the CD19 targeting structure on the surface of secreted extracellular vesicles. We systematically evaluated the therapeutic efficacy and safety of these vesicles in an MRL/lpr mouse model, aiming to provide a novel “cell-free therapy” strategy for SLE and other autoimmune diseases.

## 2. Materials and Methods

### 2.1. Cell Culture and Reagents

The human NK cell line NK-92 was generously provided by Professor Li Jing from Ocean University of China. NK-92 cells were cultured in α-MEM (Genscript, Nanjing, China) supplemented with 12.5% fetal bovine serum, 12.5% horse serum, 1% penicillin (100 U/mL) and streptomycin (100 μg/mL) (P/S), and 200 U/mL recombinant human interleukin-2 (IL-2). Human embryonic kidney (HEK293T) cells were kindly provided by Professor Gu Yuchao from Qingdao University of Science and Technology and cultured in high-glucose Dulbecco’s Modified Eagle Medium supplemented with 10% fetal bovine serum (FBS), 1% penicillin (100 U/mL) and streptomycin (100 μg/mL) (P/S) (Beyotime Biotechnology). All cell lines were cultured at 37°C in a 5% CO incubator. All FBS mentioned above was purchased from ExCell Bio (Grand Island). Antibodies were sourced as specified in the flow cytometry section.

### 2.2. Plasmid Construction, Preparation of CD19scFv-LAMP-2B Viral Particles, and Cell Transfection

First, gene sequences for Lysosome-Associated Membrane Glycoprotein 2 (LAMP-2B) and the anti-CD19 protein recognition region (CD19 scFv) were obtained from the NCBI GenBank. The target DNA fragment was synthesized and cloned into the pLVX-IRES vector backbone carrying a GFP tag using XhoI and SpeI restriction enzymes to generate the CD19-scFv expression construct; the pLVX-IRES plasmid without CD19-scFv served as a control to verify the specific therapeutic effect of CD19-scFv. HEK-293T cells in the logarithmic growth phase were seeded in 10 cm culture dishes and cultured in DMEM with 10% FBS until reaching 70%–80% confluence. The target vector plasmid was co-transfected with the helper packaging plasmid psPAX2 and the envelope plasmid pMD.2G into cells using Lipofectamine 3000 (Thermo Fisher, USA) to produce viral particles. Subsequently, NK-92 cells were transfected with the lentiviral particles via polybrene-mediated transduction to integrate CD19scFv-LAMP-2B into the cellular genome and establish the CD19scFv-NK92 cell line. Green fluorescence was observable under a fluorescence microscope 48 hours post-transfection. Cells were passaged at least 5 times under the selective pressure of 600 μg/mL G418 geneticin to achieve selection of a stably transfected CD19scFv-NK92 cell line.

### 2.3. Isolation and Characterization of Extracellular Vesicles

V-CD19-Exo were secreted by the NK-92 cell line stably expressing the CD19scFv-LAMP2B fusion protein. This cell line was routinely cultured and passaged under standard conditions (37°C, 5% CO ) using complete α-MEM medium supplemented with 12.5% fetal bovine serum, 12.5% horse serum, 1% penicillin (100 U/mL) and streptomycin (100 μg/mL) (P/S), and 200 U/mL recombinant human interleukin-2 (IL-2).

To collect engineered extracellular vesicles, cells at approximately 80% confluence were switched to serum-free medium and cultured for an additional 48 hours. The conditioned medium was collected and subjected to differential centrifugation at 4°C: first at 300×g for 10 minutes to remove live cells, then at 2000×g for 15 minutes to remove dead cells and large debris, and finally at 14,000×g for 30 minutes to remove organelles and large vesicles. The supernatant was filtered through a 0.22 μm PES membrane and initially concentrated using 50 kDa molecular weight cut-off centrifugal filter units. The concentrated solution was mixed with Exosome Precipitation Solution at a 4:1 volume ratio, incubated overnight at 4°C, and then centrifuged at 12,000×g for 30 minutes at 4°C. The pellet was resuspended in 200 μL Solution A, centrifuged again at 12,000×g for 5 minutes at 4°C, and the supernatant (containing vesicles) was collected, aliquoted, and stored at −80°C.

Particle size distribution and concentration of the obtained extracellular vesicles were determined using a Marvin NTA NS300. Typical cup-shaped morphology was observed via transmission electron microscopy. Expression of exosomal marker proteins CD9, CD63, and CD81 was detected by Western Blot to confirm vesicle purity.

### 2.4. In Vivo Treatment

Sixteen-week-old female MRL/lpr mice and sixteen-week-old female C57BL/6 mice were purchased from Beijing Vital River Laboratory Animal Technology Co., Ltd. Animals were housed in the specific pathogen-free facility of the Animal Experiment Center at Qingdao University of Science and Technology. Experiments were conducted following the National Institutes of Health Guide for the Care and Use of Laboratory Animals and were approved by the Ethics Committee of Qingdao University of Science and Technology. Sixteen-week-old MRL/lpr mice, serving as the SLE model, were randomly divided into a normal saline (NC) group (n=5), an NK92-Exo group (n=5), and a V-CD19-Exo group (n=5). Sixteen-week-old female C57BL/6 mice served as normal controls (n=5). Mice were treated with normal saline, NK92-Exo (10 particles), or V-CD19-Exo (10 particles), respectively. Mice were treated twice weekly for 3 weeks. During the treatment period, urine and serum were collected twice weekly for subsequent detection of relevant indicators. Urine was collected using the “bladder massage” method, and serum was separated from blood collected via orbital sinus. For survival studies, the same interventions and groupings (saline group, NK92-Exo group, V-CD19-Exo group) were applied, and mouse survival was observed for as long as possible. All interventions in this study, including normal saline, NK92-Exo, and V-CD19-Exo, were administered via intraperitoneal injection. Intervention details are shown in Fig. 3A. By week 19, treatment ended, mice were euthanized, and serum, kidneys, and spleens were collected from all mice. At the end of the experiment, euthanasia was performed using sodium pentobarbital according to the AVMA Guidelines for the Euthanasia of Animals (2020 Edition). Animal carcasses were disposed of following the regulations of the National Institutes of Health Guide for the Care and Use of Laboratory Animals.

### 2.5. Flow Cytometry Analysis

Mouse spleen leukocytes were isolated and prepared using a mouse spleen lymphocyte isolation kit (Tianjin Haoyang Technology Co., Ltd.) strictly following the manufacturer’s instructions to obtain a highly viable single-cell suspension. After counting cells using a hemocytometer, cells were resuspended in PBS containing 1% (w/v) bovine serum albumin (BSA, Gibco) and incubated at room temperature for 15 minutes to block non-specific binding sites. 1×10 cells were incubated with FITC-conjugated anti-mouse CD19 antibody and APC-conjugated anti-mouse CD20 antibody (purchased from BioLegend) for 30 minutes at 4°C in the dark. Cells were then washed twice with PBS containing 1% BSA and finally resuspended in 300 μL PBS buffer for analysis. Flow cytometry data acquisition was performed using a CytoFLEX flow cytometer (Beckman Coulter, USA). Subsequent data analysis was conducted using FlowJo software (Tree Star, USA).

### 2.6. Western Blot Analysis

Samples were mixed with loading buffer at a 5:1 ratio and heated at 95°C for 5 minutes in a metal bath. The treated samples were loaded onto 10% Bis-Tris SDS-PAGE gels and run sequentially in electrophoresis buffer at 80 V for 15 minutes, then 120 V for 1 hour.

Subsequently, the gel was removed, and a transfer sandwich was assembled in the order: black clamp – sponge – filter paper – protein gel – PVDF membrane – filter paper – sponge – red clamp. The transfer cassette was placed into a transfer tank immersed in ice water, and transfer was performed at a constant current of 250 mA for 2 hours. After electrophoresis, the PVDF membrane was incubated with 5% non-fat milk on a shaker (60–70 rpm) at 4°C for 2 hours to block non-specific sites. After blocking, membranes were incubated with primary antibodies against CD9, CD63, and CD81 overnight at 4°C with shaking (60–70 rpm). Membranes were then washed with 1×TBST for 10 minutes, repeated 3 times. HRP-conjugated secondary antibodies were added and incubated at room temperature for 2 hours. After incubation, membranes were washed 3 times with 1×TBST, each for 10 minutes. Immunoreactive bands were visualized using enhanced chemiluminescence (ECL, Thermo Fisher Scientific, Cat#WP20005) and imaged using a Tanon imaging system.

### 2.7. Cytokine Detection

After blood collection from the mouse orbital venous plexus, whole blood samples were allowed to clot by standing at room temperature for 2 hours. Samples were then centrifuged at 3000 rpm for 30 minutes at 4°C, and the supernatant (serum) was carefully collected. Cytokine levels were measured using ELISA kits (FANKEW) strictly following the manufacturer’s instructions. Levels of anti-double-stranded DNA antibody (anti-dsDNA), anti-nuclear antibody (ANA), immunoglobulin E (IgE), interleukin-17A (IL-17A), and interferon-γ (IFN-γ) in mouse serum were detected. All samples were run in duplicate, and concentrations were calculated based on standard curves.

### 2.8. Histological Examination

Mouse kidney tissues were fixed in formalin solution for 2 days, then dehydrated and embedded. Sections with a thickness of 4 μm were prepared. Hematoxylin and eosin (HE) staining was performed to observe pathological changes. Images were captured using a light microscope. A semi-quantitative scoring system was used to assess basic pathological changes including congestion, blood stasis, hemorrhage, edema, degeneration, necrosis, proliferation, fibrosis, tissue organization, granulation tissue formation, and inflammatory changes.

Glomerular injury was scored according to an established glomerular scoring system by evaluating 50 glomeruli and taking the average. Specific scoring categories were as follows: normal (0 points), cell proliferation or infiltration (1 point), membranoproliferation, hyaline deposition, or lobulation (2 points), global hyalinosis or crescent formation (3 points) [47].

### 2.9. Statistical Analysis

Statistical analysis of the data in this study was performed using GraphPad Prism 10 software. All values are expressed as mean ± standard deviation (SD). Data were assessed for normal distribution and similar variance between groups. One-way ANOVA was used for statistical analysis of multiple comparisons. For sample data not following a normal distribution, the Kruskal–Wallis test with post-hoc multiple comparisons was used. Pearson correlation coefficient was used to evaluate linear correlations between data, and Spearman correlation coefficient was used for non-linear correlation analysis to ensure accuracy and reliability of results. *p < 0.05, **p < 0.01, ***p < 0.001, ****p < 0.0001, with p < 0.05 considered statistically significant.

## 3. Results

### 3.1. Preparation of Engineered NK-92 Extracellular Vesicles Expressing Anti-CD19 Chimeric Antigen Receptor Protein

To endow NK-92 cell-derived extracellular vesicles with B-cell targeting capability, we employed an engineering strategy to modify the vesicle surface. The gene encoding the CD19 single-chain antibody fragment (scFv) was linked to the N-terminus of the vesicle surface marker protein LAMP-2B, fused with the GFP gene sequence, and cloned into the pLVX-IRES vector (Fig. 1A). Recombinant lentiviral particles were produced by transfecting HEK293T cells.

**Figure 1.**
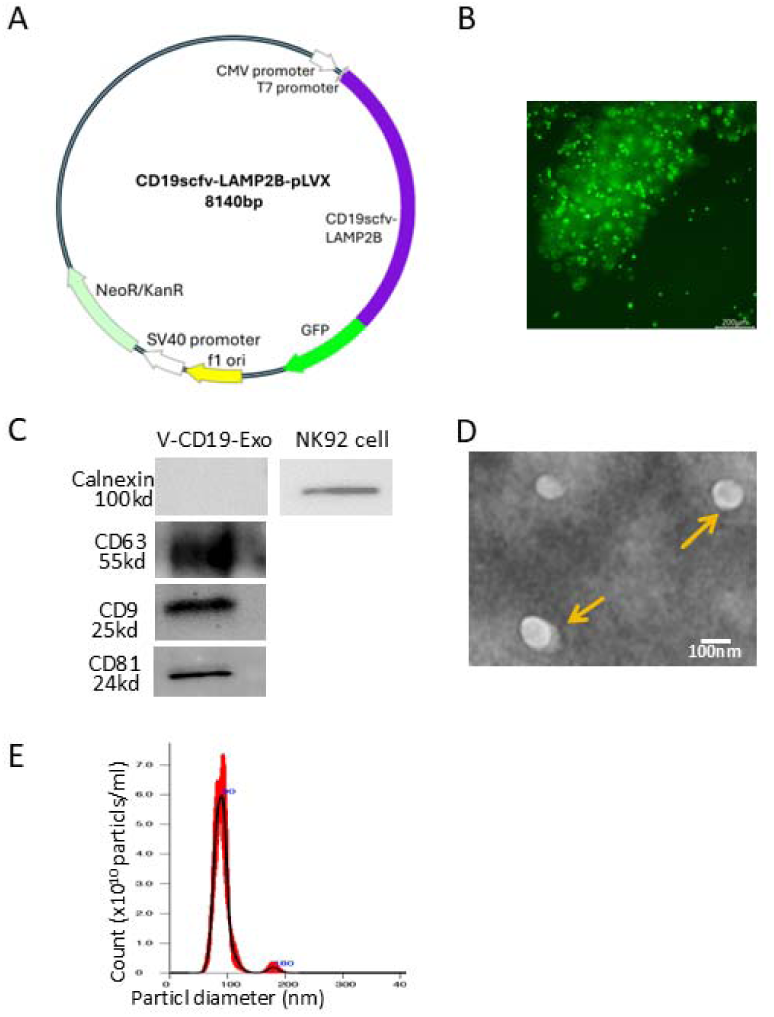
Construction and characterization of CD19-targeting NK-92-derived extracellular vesicles. (A) Construction of the CD19-targeting CD19scFv-LAMP2B-pLVX plasmid (CD19 plasmid). The CD19-targeting receptor gene was generated by fusing an anti-CD19 single-chain variable fragment (scFv) to the N-terminus of the exosomal membrane protein LAMP2B and linking it to the N-terminus of GFP. The fusion gene sequence was cloned into the multiple cloning site of the plasmid. (B) Fluorescence microscopy image of NK-92 cells 72 hours after transfection with the lentivirally packaged CD19 plasmid (V-CD19). Scale bar = 200 µm. (C) Western blot analysis of exosomal markers in V-CD19-Exo. The exosomes positively expressed CD63 (55 kDa), CD9 (25 kDa), and CD81 (24 kDa), while the endoplasmic reticulum protein Calnexin (100 kDa) was negative. (D) Transmission electron microscopy (TEM) image of extracellular vesicles (V-CD19-Exo) isolated from the supernatant of NK-92 cells after V-CD19 plasmid transfection, negatively stained. Arrows indicate cup-shaped vesicles. Scale bar = 100 nm. (E) Nanoparticle tracking analysis (NTA) of the particle size distribution and concentration of V-CD19-Exo. The exosomes exhibited a unimodal size distribution with a peak diameter of approximately 100 nm and a corresponding concentration of approximately 6.0×10¹□ particles/mL.

Subsequently, NK92 cells were transduced with the prepared lentivirus. Following antibiotic selection, an engineered cell line stably expressing the CD19scFv-LAMP-2B fusion protein was established. Observation under a fluorescence microscope with a blue light source revealed abundant green fluorescence within cell clusters, approximately 60%, confirming successful expression of CD19scFv-LAMP-2B-GFP (Fig. 1B). Subsequently, extracellular vesicles (V-CD19-Exo) were isolated from the supernatant of the CD19scFv-NK-92 cell line using an exosome extraction kit (precipitation method). To comprehensively verify the successful isolation of intact V-CD19-Exo, we characterized the obtained vesicles using Western Blot, NTA, and TEM. Western Blot results showed high expression of the exosomal marker proteins CD63, CD9, and CD81 in the obtained vesicle preparation. The endoplasmic reticulum resident protein

Calnexin was present only in the positive control parent cell lysate and was completely absent in the extracellular vesicle sample. This result confirmed the high purity of our extracellular vesicle preparation, essentially excluding contamination by cell debris or organellar components, and indicated that CD19scFv-LAMP-2B-GFP was successfully incorporated into the vesicles.

Nanoparticle tracking analysis showed that the vesicle concentration was adjusted to 1.0 × 10¹ particles/mL, with a highly concentrated particle size distribution; over 90% of particles had diameters between 50 and 150 nm (Fig. 1D), consistent with the classical size range of exosomes. Finally, transmission electron microscopy revealed that V-CD19-Exo exhibited a cup-shaped membranous vesicle morphology (Fig. 1E), characteristic of typical exosomes. In summary, the combined results from Western Blot, NTA analysis, and TEM observation collectively demonstrated the successful acquisition of engineered V-CD19-Exo vesicles.

These results collectively indicate that we successfully constructed an NK-92 cell line stably expressing CD19scFv and prepared engineered extracellular vesicles (V-CD19-Exo) with defined targeting epitopes, laying the material foundation for subsequent functional and efficacy studies.

### 3.2. V-CD19-Exo Specifically Kill Tumor Cells and Target CD19□ B Cells In Vivo

To verify whether the engineered extracellular vesicles retained the effector functions of their parent NK-92 cells after modification and to assess their potential impact on non-target cells, we first evaluated the in vitro cytotoxicity of V-CD19-Exo. They were co-cultured with tumor cells (A549) and normal mesenchymal stem cells (MSCs) for 24 hours. Assessment was performed via real-time microscopy, trypan blue exclusion counting, and cell morphological analysis. Results showed that V-CD19-Exo exhibited strong killing activity against A549 cells. After 24 hours of co-culture, microscopy revealed numerous A549 cells rounding up, detaching, and floating in the medium, with a marked reduction in adherent cell numbers (Fig. 2A), achieving a cell death rate of approximately 80%. In contrast, under identical treatment conditions, the growth status of normal MSCs was not significantly affected. Microscopy showed MSCs retained their typical spindle-shaped adherent morphology, with cell density comparable to the control group. This indicates that the engineered V-CD19-Exo successfully preserved the cytotoxicity of NK-92 cells and exhibited selective killing action, and the engineering process did not affect their cytotoxicity. Subsequently, we further validated the in vivo targeting efficacy of these vesicles in the MRL/lpr mouse model. MRL/lpr mice were randomly grouped and subjected to systemic administration for three weeks. V-CD19-Exo was administered via intraperitoneal injection at 1.0 × 10 particles/mL based on body weight. After treatment, mice were euthanized, splenic lymphocytes from each group were isolated, and changes in B cell subsets were analyzed by flow cytometry. Single-cell suspensions were stained with anti-CD19 and anti-CD20 (B-cell specific marker) antibodies to detect the proportion of CD19 CD20 double-positive mature B cells (Fig. 2B, C). Results showed that the percentage of CD19 CD20 double-positive mature B cells in the spleens of mice from the V-CD19-Exo treatment group was 1.51%, significantly lower than that in the PBS control group (10.53%, p < 0.0001) and the NK92-Exo control group (8.03%, p < 0.0001) (Fig. 2C), both showing highly significant decreases. This result directly demonstrates that V-CD19-Exo vesicles, via the CD19scFv targeting structure on their surface, can effectively home to and act on abnormally activated CD19 B cells in vivo. This finding aligns with recent research directions utilizing engineered extracellular vesicles to target B cells. For instance, similar studies have confirmed that vesicles displaying anti-CD19 single-chain antibodies can specifically recognize and eliminate CD19 B cells in vitro and in vivo [48].

**Figure 2.**
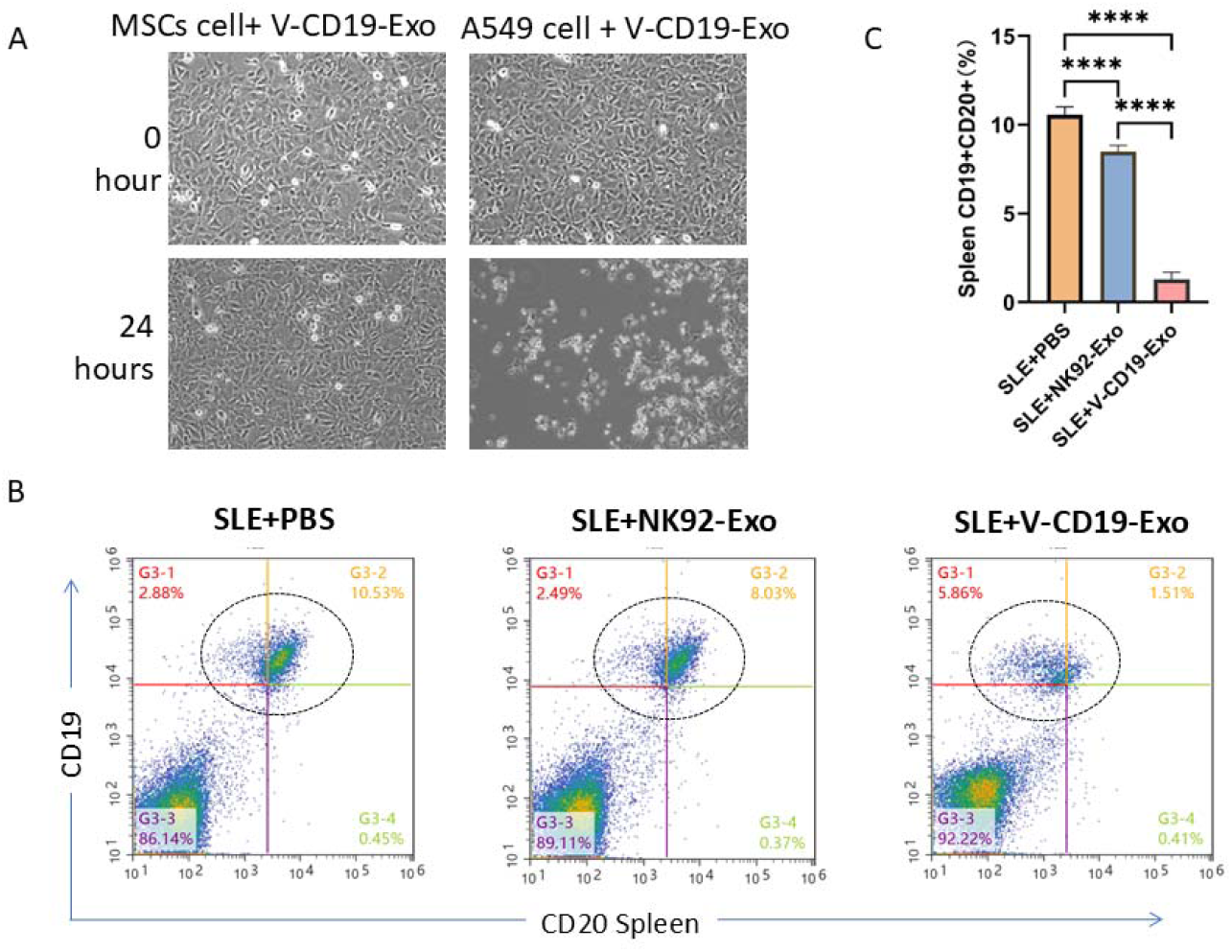
V-CD19-Exo retains NK-92 cytotoxicity against tumor cells and selectively targets CD19□ B cells in vivo. (A) Cytotoxic effect of V-CD19-Exo on A549 tumor cells. Tumor cells were seeded in 6-well plates containing DMEM + 10% FBS (1x10□ A549 cells/2 mL/well), cultured for 24 hours, then V-CD19-Exo was added, mixed, and cultured for another 24 hours. (B) Effect of V-CD19-Exo on B cells in MRL/lpr mice. 16-week-old female MRL/lpr mice were intraperitoneally injected with V-CD19-Exo (10□ particles/mouse/time). Mice were euthanized 3 weeks later, spleens were collected to prepare single-cell suspensions, and splenic B cell populations were analyzed by flow cytometry using anti-CD20-FITC and anti-CD19-APC antibodies. NC, control group; NK92-EXO, MRL/lpr mice injected with NK92 cell exosomes; CD19scfv-NK92-Exos, MRL/lpr mice injected with V-CD19-Exo exosomes. The dotted box indicates CD20 and CD19 double-positive B cells. (C) Quantitative analysis of CD20 and CD19 double-positive B cells (cells within the circular gate in Figure B) in the spleens from mice treated with NC, control, and NK92-EXO groups, corresponding to Figure B.

In summary, V-CD19-Exo successfully integrated the inherent cytotoxicity of NK-92 vesicles with the targeting ability mediated by CD19scFv, achieving specific action against CD19 B cells and resulting in a significant reduction of mature B cell populations in the spleen. This provides a basis for its potential systemic therapeutic effect in SLE models and offers key functional evidence for its application in treating B cell-related diseases.

### 3.3. V-CD19-Exo Effectively Alleviates Lupus Nephritis and Renal Inflammation in MRL/lpr Mice

#### 3.3.1. Evaluation of the Renal Protective Effect of V-CD19-Exo in SLE Model Mice

To assess the renal protective effect of V-CD19-Exo in systemic lupus erythematosus (SLE) model mice (Fig. 3A), we systematically analyzed renal function and pathological changes in MRL/lpr mice. Dynamic monitoring showed that urinary protein levels in untreated MRL/lpr mice continuously increased with disease progression, reaching ∼2000 mg/L by the end of the experiment, indicating severe proteinuria, a hallmark of lupus nephritis. In contrast, following V-CD19-Exo treatment, the upward trend of urinary protein was significantly delayed. Throughout the observation period, urinary protein levels in the treatment group remained consistently low, averaging below 1000 mg/L by the end of the experiment, showing a statistically significant difference compared to the NC group (Fig. 3C). Concurrently, the urinary protein/creatinine ratio (UPCR), a more stable indicator of renal function, was also significantly reduced in the V-CD19-Exo treatment group (Fig. 3D). Vesicles obtained from NK-92 cells transfected with the control plasmid did not show this significant therapeutic effect. These results, consistent with the trend in urinary protein content, confirm that V-CD19-Exo treatment effectively reduces protein leakage in MRL/lpr mice, ameliorates the glomerular filtration barrier function, and indicates protection of renal function.

**Figure 3.**
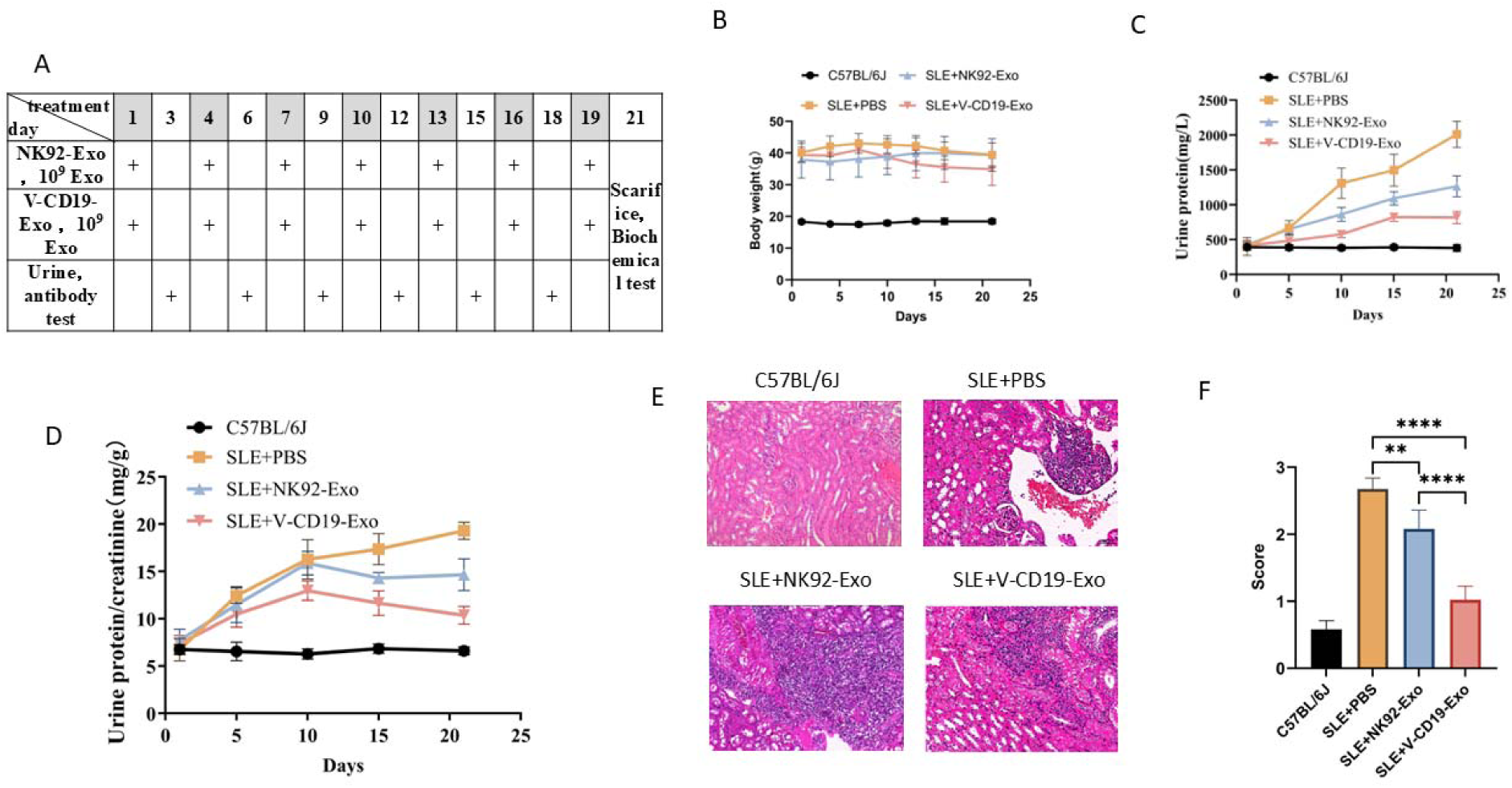
Effects of V-CD19-Exo on body weight, proteinuria, and renal histopathology in MRL/lpr mice. Treatment and detection timelines are as indicated. (A) Drugs were injected according to the experimental design timeline, or mouse urine and serum were collected to monitor urinary protein, urinary creatinine, and serum autoantibody levels. Mice were euthanized at the experimental endpoint, and kidneys were collected for pathological analysis. Experimental data analysis is shown as follows: (B) Dynamic changes in mean body weight of mice in each group, (C) Kidney function indicator - urinary protein concentration, (D) Urinary protein to urinary creatinine ratio. Data presented as mean ± SEM. **p < 0.01, ***p < 0.001 using two-way ANOVA analysis. (E) Representative images of renal histopathology at the experimental endpoint (H&E staining, scale bar = 100 μm), showing glomerular necrosis (black arrows) and lymphocyte infiltration (white arrows). (F) Kidney injury score, **p<0.001, n=5; **** p<0.0001, n=5.

Hematoxylin-eosin (HE) staining of kidney tissue was performed to evaluate the effect of V-CD19-Exo on renal pathological changes in MRL/lpr mice. Histological analysis revealed that kidneys from mice in the NC group exhibited typical active lupus nephritis lesions, including diffuse glomerular lymphocyte infiltration, mesangial matrix proliferation, accompanied by tubular epithelial cell swelling and interstitial edema (Fig. 3E). In contrast, renal histopathological damage was significantly attenuated in the V-CD19-Exo treatment group.

Inflammatory cell infiltrates in the glomerular area were markedly reduced, tubular structures were largely intact, and interstitial edema was improved (Fig. 3E). Subsequent evaluation of glomerular lesions using a semi-quantitative histological scoring system showed that the total renal pathology score in the V-CD19-Exo treatment group was significantly lower than that in the NC group (Fig. 3F, p < 0.001, n=5; p < 0.0001, n=5). These results indicate that V-CD19-Exo treatment effectively inhibits local lymphocyte infiltration and activation in the kidney, reduces the inflammatory response mediated by immune complex deposition, and thereby significantly ameliorates renal pathological damage in SLE model mice.

The above results demonstrate that V-CD19-Exo treatment, while targeting autoreactive B cells, can alleviate the progression of lupus nephritis at the pathological structural level, exhibiting excellent renal protective effects.

#### 3.3.2. Unveiling the Systemic Immunomodulatory Effects of V-CD19-Exo

To investigate whether V-CD19-Exo exerts systemic immunomodulatory effects, we evaluated its impact on spleen weight, serum cytokine levels (IgE, IL-17A, and IFN-γ), and anti-double-stranded DNA IgG autoantibody levels in MRL/lpr mice. The results showed that the treatment effectively reversed SLE-associated abnormal immune states at multiple levels. First, we observed that V-CD19-Exo treatment significantly ameliorated the pathological phenotype of immune organs in model mice. Compared to the PBS control group, which exhibited significant splenomegaly, V-CD19-Exo treatment restored spleen size and weight to near-normal levels (Fig. 4A, B). Pathological splenomegaly, a direct indicator of systemic immune hyperactivation in MRL/lpr mice, was thus reversed. This result indicates that V-CD19-Exo treatment significantly ameliorated systemic immune inflammation in model mice. Simultaneously, we used ELISA to measure serum levels of IgE, IL-17A, and IFN-γ, finding that V-CD19-Exo treatment significantly downregulated these pro-inflammatory factors. Compared with the NC group and the control NK92-Exo treatment group, V-CD19-Exo treatment significantly reduced serum levels of IgE, IL-17A, and IFN-γ in MRL/lpr mice (Fig. 4C–E). This change reflects an overall improvement in the body’s immune inflammatory status.

**Figure 4.**
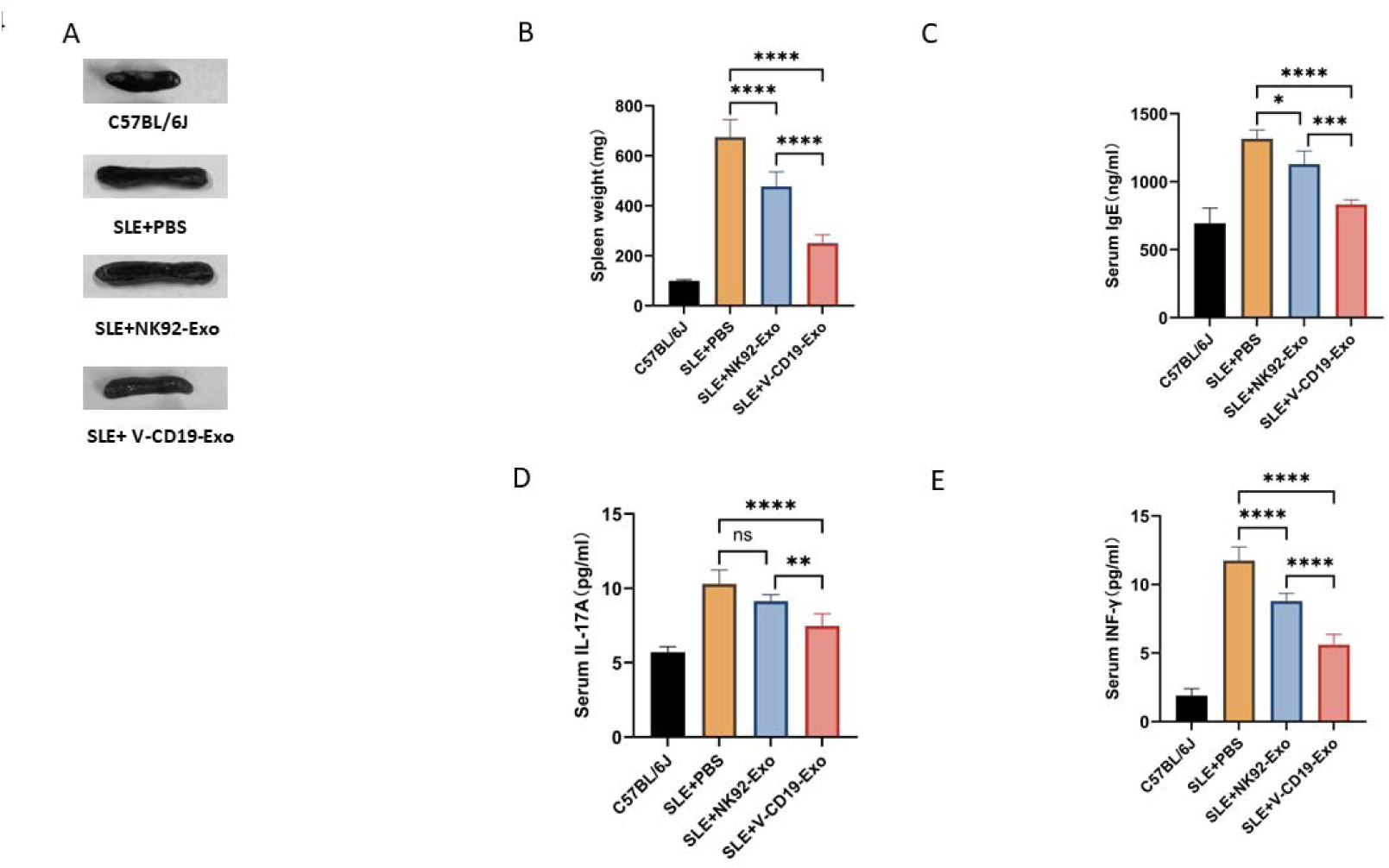
V-CD19-Exo treatment alleviates splenomegaly and systemic inflammation in MRL/lpr mice. (A) Representative images of spleens at the treatment endpoint (week 19). Spleens from healthy control C57BL/6J mice, untreated MRL/lpr mice, control NK92-Exo treatment group, and V-CD19-Exo treatment group. Scale bar: 1 cm. (B-E) Quantitative analysis of spleen weight and serum inflammatory cytokines at the treatment endpoint. (B) Spleen weight. (C) Serum total IgE concentration. (D) Serum IL-17A concentration. (E) Serum IFN-γ concentration. Data are presented as mean ± standard error of the mean (n=X per group). Statistical analysis was performed using one-way ANOVA with Tukey’s post-hoc test. *p < 0.05, **p < 0.01, ***p < 0.001; ns, not statistically significant.

More critically, serological analysis showed that V-CD19-Exo treatment significantly reduced the titers of two key SLE autoantibodies—anti-double-stranded DNA (dsDNA) antibody and anti-nuclear antibody (ANA)—showing a downward trend (Fig. 5A, B). This effect was significantly superior to the non-targeted NK92-Exo control group, demonstrating that CD19scFv-mediated targeted delivery offers a specific advantage in inhibiting autoantibody production. In summary, V-CD19-Exo achieved multi-level systemic immunomodulation by restoring immune organ homeostasis, downregulating broad-spectrum pro-inflammatory cytokines, and specifically inhibiting pathogenic autoantibody production.

**Figure 5.**
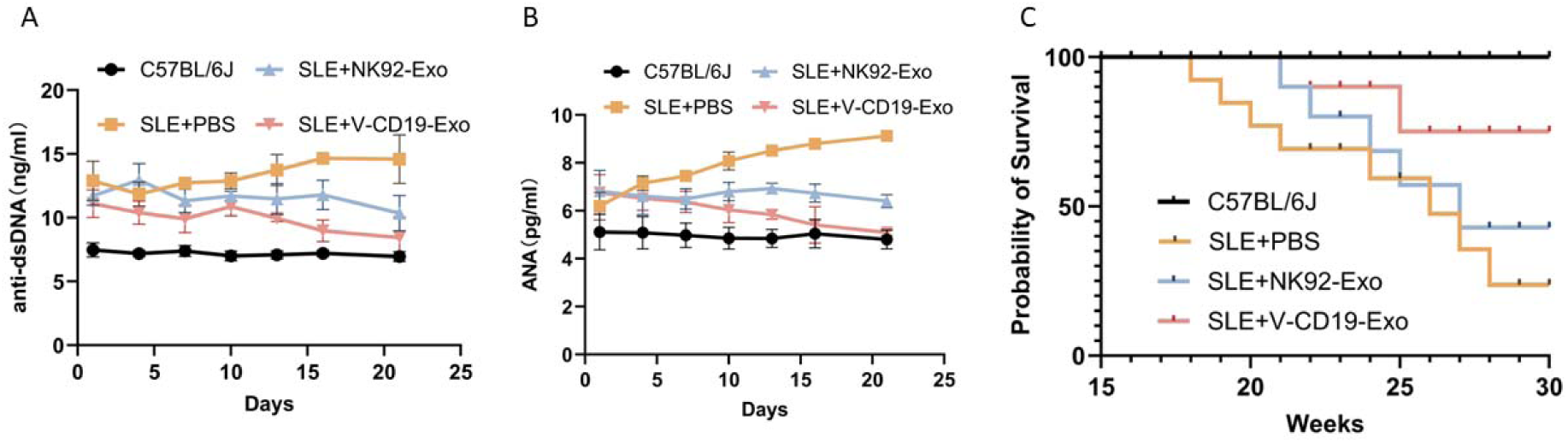
Autoantibody levels and survival during V-CD19-Exo treatment in MRL/lpr mice. (A) Serum anti-double-stranded DNA IgG antibody levels, and (B) Anti-nuclear antibody titers (n=5). (C) Kaplan-Meier survival curve of MRL/lpr mice over a 30-week observation period (n=10).

#### 3.3.3. Does Immune Improvement Translate into Long-Term Survival Benefits?

Survival rate is the most direct and critical indicator for evaluating SLE treatment efficacy. We monitored the survival of mice in each group for up to 30 weeks, revealing significant differences in survival among the different treatment groups. Mice in the NC group began to die from week 17, with a survival rate of only 25% by the experimental endpoint. The survival curve of the control NK92-Exo group was similar. In stark contrast, the survival rate of mice in the V-CD19-Exo treatment group was significantly improved, reaching approximately 80% (Fig. 5C), indicating that this treatment significantly prolongs the lifespan of model mice.

In summary, long-term survival observations, together with the previously observed renal function protection and reduced pathological damage, strongly demonstrate that V-CD19-Exo treatment not only effectively regulates autoimmune activity but also fundamentally and significantly prolongs the survival of disease model animals, demonstrating clear therapeutic potential and protective effects.

## 4. Discussion

Natural killer (NK) cells are a crucial component of the innate immune system, capable of specifically killing target cells without prior antigen presentation. NK cells and their derived extracellular vesicles play important roles in tumor immunotherapy, maintaining immune homeostasis, and regulating aberrant immune responses.

NK-92 cells are an immortalized human NK cell line derived from a lymphoma patient and are widely used in tumor immunotherapy research. In recent years, extracellular vesicles derived from NK-92 cells have gained increasing attention. They are considered to inherit, to some extent, the immunomodulatory and targeted killing functions of the parent cells while avoiding the safety concerns associated with live cell therapy. However, no reports have yet been published on the application of NK-92-derived extracellular vesicles in the treatment of autoimmune diseases.

This study demonstrates for the first time that engineered NK-92 cell-derived extracellular vesicles can significantly reduce the number of CD19 B cells in vivo. In contrast, vesicles produced by NK-92 cells transfected with a control plasmid showed no significant therapeutic effect, suggesting that vesicles targeting CD19 B cells can effectively regulate abnormal B cell activation or proliferation. This finding provides a novel cell-free therapeutic strategy for B cell-mediated autoimmune diseases such as rheumatoid arthritis and systemic lupus erythematosus, significantly expanding the application scope of NK cell therapy.

This study successfully constructed engineered extracellular vesicles (V-CD19-Exo) derived from NK-92 cells, conferring targeting ability via CD19scFv expression, and confirmed their significant therapeutic efficacy in the MRL/lpr lupus mouse model. This strategy effectively alleviated renal pathology, systemic immune dysregulation, and autoantibody production, ultimately prolonging survival. These results indicate that targeted therapeutic strategies based on engineered extracellular vesicles offer a new interventional approach for SLE.

Compared to cell therapy, extracellular vesicles offer distinct advantages. NK cell culture is time-consuming; in vitro established NK cell lines after plasmid transfection may exhibit reduced cytotoxicity, limited passage numbers, and risks like cytokine storms and immunogenicity. NK-92 cells, as an immortalized cell line, carry potential tumorigenicity risks and require irradiation before use, which affects their in vivo survival time and therapeutic persistence. Both cell types lack crucial homing receptors, limiting their ability to target specific cells or tissues. In contrast, targeted modified NK-92 vesicles lack proliferative capacity, possess low immunogenicity, offer higher safety, and are amenable to engineering modifications and large-scale production. Therefore, this approach completely eliminates dependence on live cell infusion, avoiding safety risks associated with cell therapy. Simultaneously, based on existing cell culture and vesicle isolation technologies, this platform holds potential for scalable production, cost control, and high batch-to-batch consistency, providing a novel and translationally feasible solution for targeted therapy of autoimmune diseases with promising clinical prospects.

This study established a modular engineered extracellular vesicle delivery platform. By anchoring the targeting molecule CD19 single-chain antibody fragment (CD19 scFv) onto the vesicle membrane surface, the vesicles were endowed with active recognition, specific binding, and homing capabilities towards CD19 B cells, achieving precise targeted delivery. This design enables vesicles to efficiently act on pathogenic B cells, providing a new technological pathway for targeted intervention in B cell-related diseases. Through directed engineering of EVs, their yield, targeting ability, and therapeutic specificity can be systematically enhanced, thereby overcoming the limitations of natural EVs [38, 39]. The CD19 scFv “targeting module” can be flexibly replaced with other targeting molecules (e.g., scFvs against different tumor antigens or immune cell markers) depending on the disease type. Concurrently, the internal cavity of the vesicle can be loaded with various therapeutic payloads, including small molecule drugs, nucleic acids, or functional proteins. This forms a multifunctional delivery system with a “replaceable target head and variable payload,” granting the platform broad disease adaptability and good industrialization potential [19, 49, 50].

The core innovation of this study lies in adapting the CAR targeting design concept, replacing immune cell therapy with extracellular vesicles. By constructing a stable cell line, continuously and stably produced targeted extracellular vesicles are applied for treatment, offering a novel solution for precise intervention in autoimmune diseases and B cell-related disorders.

Despite the significant potential shown by engineered NK-92 vesicles in targeted therapy, their clinical translation still faces multiple challenges. Large-scale, high-purity production of extracellular vesicles remains a critical bottleneck for industrialization. Current yields from cell culture systems are limited, and preparations are prone to contamination with other extracellular vesicles or impurities, potentially affecting product uniformity and therapeutic efficacy.

This study clearly demonstrates the effectiveness of targeting B cells therapeutically. However, the precise cellular mechanism of action warrants further investigation. Firstly, whether the vesicle-mediated reduction in B cells results from direct cell elimination (killing) or downregulation of CD19 gene activity remains unclear. This study observed a sharp decline in target cell proportion in the spleen (from 10.53% to 1.51%) accompanied by improvements in systemic immune parameters, suggesting a potential direct cytotoxic effect, given that the plasmid itself lacks the function to recognize the CD19 gene. The observed outcome is likely mediated by cytotoxic proteins (e.g., perforin, granzymes) carried by NK-92 cell-derived vesicles.

Notably, although within the same MRL/lpr model, we observed that identical treatment exhibited more significant efficacy in male mice compared to female mice (data not shown). This could stem from inherent sex differences in the model—females display earlier onset and more robust autoimmune phenotypes, making the same intervention relatively more effective in males with lower inflammatory baselines. Alternatively, it suggests that differences in the immune microenvironment of target organs like the kidney might influence EV efficacy, necessitating deeper mechanistic exploration considering sex as a biological variable in future studies.

## Acknowledgements

Competing interests The authors declare that they have no competing interests.Funding This study was supported by the Qingdao Natural Science Foundation (Grant No. 33122025003).

## CRediT Author Statement

**Yu Sun:** Conceptualization, Methodology, Investigation, Writing – original draft. **Zeyu Tang:** Investigation, Methodology. **Mengting Guo:** Investigation, Data curation. **Zimeng Zhai:** Investigation. **Zixian Wu:** Investigation. **Xia Wang:** Investigation. **Fang Li:** Investigation. **Weiling An:** Resources. **Xiaowei Dou:** Conceptualization, Supervision, Funding acquisition, Writing – review & editing.

